# NOBIAS: Analyzing anomalous diffusion in single-molecule tracks with nonparametric Bayesian inference

**DOI:** 10.1101/2021.07.15.452497

**Authors:** Ziyuan Chen, Laurent Geffroy, Julie S. Biteen

## Abstract

Single particle tracking (SPT) enables the investigation of biomolecular dynamics at a high temporal and spatial resolution in living cells, and the analysis of these SPT datasets can reveal biochemical interactions and mechanisms. Still, how to make the best use of these tracking data for a broad set of experimental conditions remains an analysis challenge in the field. Here, we develop a new SPT analysis framework: NOBIAS (<u>NO</u>nparametric <u>B</u>ayesian <u>I</u>nference for <u>A</u>nomalous Diffusion in <u>S</u>ingle-Molecule Tracking), which applies nonparametric Bayesian statistics and deep learning approaches to thoroughly analyze SPT datasets. In particular, NOBIAS handles complicated live-cell SPT data for which: the number of diffusive states is unknown, mixtures of different diffusive populations may exist within single trajectories, symmetry cannot be assumed between the *x* and *y* directions, and anomalous diffusion is possible. NOBIAS provides the number of diffusive states without manual supervision, it quantifies the dynamics and relative populations of each diffusive state, it provides the transition probabilities between states, and it assesses the anomalous diffusion behavior for each state. We validate the performance of NOBIAS with simulated datasets and apply it to the diffusion of single outer-membrane proteins in *Bacteroides thetaiotaomicron*. Furthermore, we compare NOBIAS with other SPT analysis methods and find that, in addition to these advantages, NOBIAS is robust and has high computational efficiency and is particularly advantageous due to its ability to treat experimental trajectories with asymmetry and anomalous diffusion.

## Introduction

The biophysical dynamics of biomolecules reflect the biochemical interactions in the system, and these dynamics can be quantified within a dataset of single-particle trajectories obtained by tracking individual molecules. The invention of the super-resolution microscope (Moerner and Kador, 1989; Hell and Wichmann, 1994; Betzig et al., 2006; Hess et al., 2006; Rust et al., 2006) and single-particle tracking (SPT) methods (Yildiz, 2003; Deich et al., 2004; Elmore et al., 2005; Manley et al., 2008) have made possible investigations of biomolecular dynamics at a high temporal and spatial resolution both *in vitro* and *in vivo*. Moreover, quantitative SPT algorithms can connect the real-time dynamics from biophysical trajectories to biochemical roles to uncover whether a molecule interacts with other cellular components (Izeddin et al., 2014), freely diffuses (Badrinarayanan et al., 2012), is actively transported (Park et al., 2014), or is constrained to a certain region (Bayas et al., 2018).

Conventionally, SPT trajectory datasets have been assumed to be Brownian, such that the mean squared displacement, MSD, of each track is linearly proportional to the time lag, *τ*, and the diffusion coefficient, *D*, can be calculated from a linear fit to this curve (Qian et al., 1991; Saxton, 1997). This Brownian motion assumption works accurately for freely diffusing molecules in solution. Despite the accessibility of this method, it has a simplified assumption that the molecule is freely diffusing with a single diffusive state (a single *D* value) for each trajectory. In the complicated cellular environment, however, multiple diffusive states, each characterized by an average *D*, can exist—for instance due to binding and unbinding events— and molecules can transition between different states to produce heterogeneity even within single trajectories. To reveal these heterogeneous dynamics, probability distribution-based methods such as cumulative probability distribution (Schütz et al., 1997; Mazza et al., 2012), have been applied. Probability distribution-based models use kinetic modeling with a predetermined number of diffusive states and are fit to histograms of displacements calculated at different time lags. These probability-based kinetic models pool displacements from the SPT dataset to estimate the *D* and weight fraction for each diffusive state in the model. Probability distribution-based analytical tools (Rowland and Biteen, 2017; Hansen et al., 2018) have been widely applied to SPT datasets with extra corrections that consider the experimental microscopy data collection process. These corrections include localization error (Michalet and Berglund, 2012), confinement (Kusumi et al., 1993), motion blur (Berglund, 2010; Deschout et al., 2012), and out-of-focus effects (Lindén et al., 2017) in the probability model.

For some well-studied biological systems in which the biochemical states of molecules have been determined through other methods, a fixed-state number analytical tool can be suitable for quantifying the dynamics and weight for each state (Elf et al., 2007; Hansen et al., 2017). However, SPT can also be used as the beginning step to investigate biomolecule dynamics without prior knowledge of how many diffusive states there supposed to be Monnier et al., 2015; Sungkaworn et al., 2017; Biswas et al., 2021). In these cases, how to objectively determine the number of diffusive states is a great challenge. Moreover, these models provide a *D* value for each subpopulation, but they do not assign the diffusive state to each individual single-molecule step, nor do they quantify the transition probability between distinct diffusive states within one trajectory. However, these transition probabilities can reveal important biological meaning such as the presence of critical biochemical intermediates (Biswas et al., 2021).

Bayesian statistics and Hidden Markov Models (HMMs) have been applied to analyze SPT datasets without assuming a predetermined number of diffusive states and to access the probabilities of transitioning between distinct states (Persson et al., 2013; Monnier et al., 2015; Karslake et al., 2020; Heckert et al., 2021). vbSPT, which was one of the first applications of HMM for SPT analysis (Persson et al., 2013), uses a maximum-evidence criterion to select between models with different numbers of diffusive states; within each model, a fixed-order HMM is used to infer the diffusion coefficient, weight fraction, and transition probabilities for each state. More recently, nonparametric Bayesian models based on Dirichlet processes were combined with HMM to recover the number of diffusive states from SPT trajectory datasets, such as in SMAUG (Karslake et al., 2020) and DSMM (Heckert et al., 2021). In these models, the motion of the molecule is approximated to be symmetric and Brownian, which is an oversimplification considering the crowded environment and various interaction partners for biomolecules in cells.

To move beyond Brownian motion, here we consider a more general random walk family: anomalous diffusion. In anomalous diffusion, MSD and *τ* are related by a power law distribution, *MSD*~*τ*^α^, where *α* is the anomalous diffusion exponent (Metzler et al., 2014). Brownian motion is a special case of anomalous diffusion (*α* = 1), and other cases can be further divided into subdiffusion (*α* > 1) and superdiffusion (*α* < 1). Biomolecules have been reported to diffuse anomalously in many situations, such as constrained membrane protein motion (Jeon et al., 2016), the facilitated diffusion of DNA binding protein (Bauer and Metzler, 2012), and active transportation of cargoes (Caspi et al., 2002). Different designs of neural networks effectively classify the diffusion type of trajectories (Bo et al., 2019; Granik et al., 2019; Argun et al., 2021; Gentili and Volpe, 2021), however these analyses typically assume that each track is dynamically homogeneous and is characterized by a single type of diffusion and a single *D* value. It remains a challenge to classify the diffusion type within a trajectory when considering the possibility of changes in dynamics or diffusion types within a single track.

Here we introduce the <u>NO</u>nparametric <u>B</u>ayesian Inference for <u>A</u>nomalous diffusion in <u>S</u>ingle-molecule tracking (NOBIAS) framework to address the assumptions and simplifications discussed above and provide a more physiologically relevant analysis algorithm to quantify the dynamics encoded in SPT datasets (**Figure 1**). In particular, NOBIAS recovers the number of diffusive states and predict the diffusion type for each diffusive state, even in heterogeneous trajectories. The NOBIAS framework consists of two modules. The first module uses a Hierarchical Dirichlet Process Hidden Markov Model (HDP-HMM) with multivariate Gaussian emission to recover the number of diffusive states and infer their corresponding diffusion coefficients and weight fractions. This module also assigns each single-molecule step a diffusive state label to provide the state label sequence and the matrix of transition probabilities. In the second module, the original trajectories are segmented by diffusive state label and a pre-trained Recurrent Neural Network (RNN) is used to classify these segments and assign the diffusion type (Brownian motion, Fractional Brownian motion, Continuous Time Random Walk, or Lévy Walk) for each diffusive state. We simulated trajectory datasets with mixtures of heterogeneous dynamics and diffusion types to validate the NOBIAS framework, and we analyzed the SPT dataset from experimental measurements of the SusG outer-membrane protein in living *Bacteroides thetaiotaomicron* to access its dynamics and anomalous diffusion behaviors, which are consistent with its role in starch catabolism in gut microbiome. This framework uses nonparametric Bayesian statistics and Deep learning to thoroughly analyze a single-molecule tracking dataset. It provides an objective method to determine the number of diffusive states in an SPT dataset and accesses the multidirectional dynamics of each state. A further diffusion type classification for each diffusive state is also included in the framework. The NOBIAS framework overcomes some oversimplified assumptions in SPT analysis and provides a powerful tool to fully make use of single-molecule tracking data.

**Figure 1.**
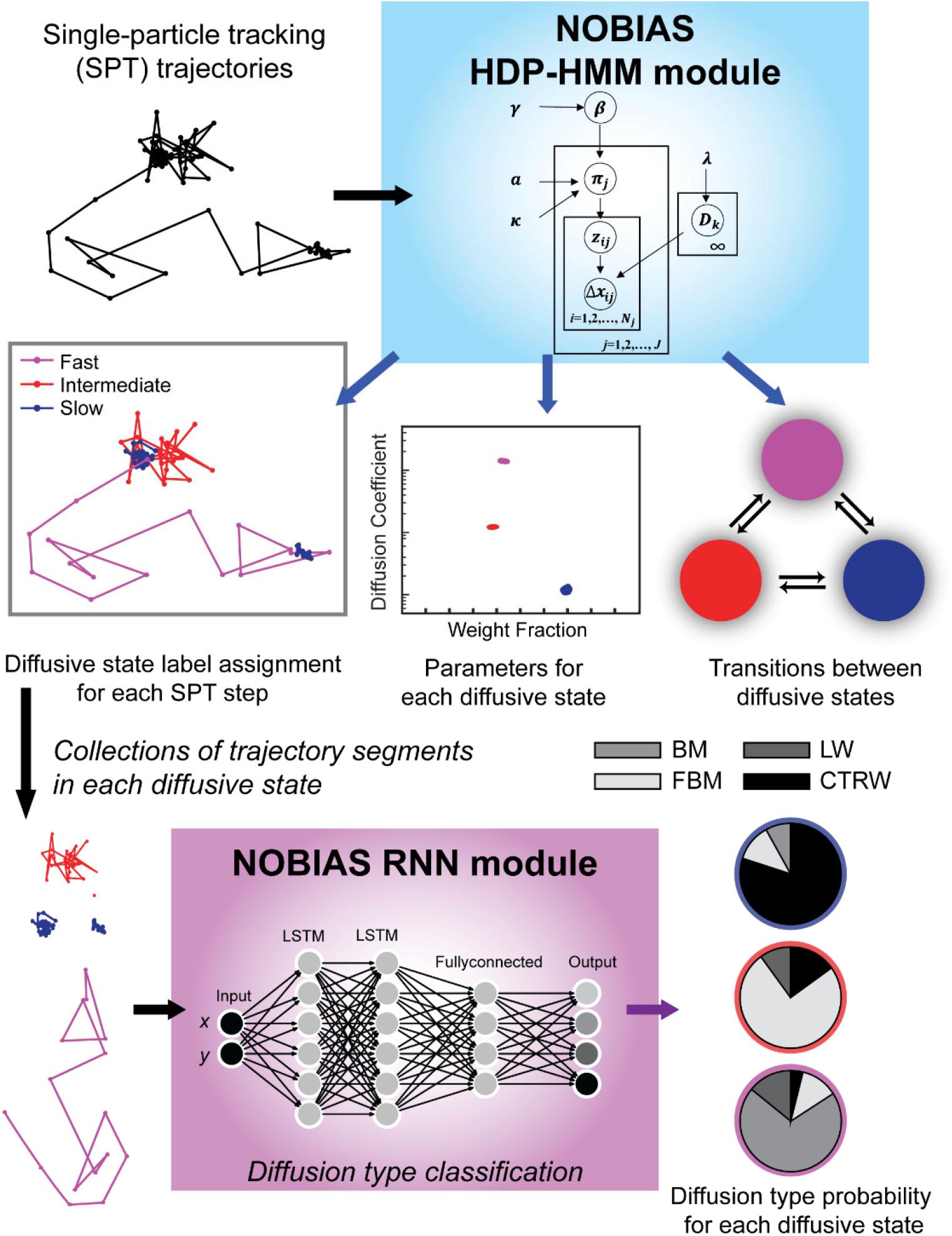
NOBIAS workflow. (1) Single-particle tracking (SPT) trajectory datasets are processed in the NOBIAS HDP-HMM module: the observed data (the displacements, Δ***x***) are analyzed in the context of the emission parameters (the diffusion coefficients, ***D***). The state sequence, *z*, indicates the diffusive state corresponding to each step, and the transition matrix, ***π***, is estimated with a Hierarchical Dirichlet process prior using concentration hyperparameters *a* and *γ* and the sticky parameter, *κ*. The HDP-HMM module provides *D* and the weight fraction for each diffusive state, the π for transition probabilities between these states, and a state label assignment for each SPT step. (2) In the NOBIAS RNN module, trajectory segments of the same diffusive state are collected and put in a pre-trained Recurrent Neural Network (RNN) with two long short-term memory (LSTM) layers to classify the diffusion type for each diffusive state.

## Methods

### Hidden Markov model (HMM)

A HMM infers a system with a discrete-valued sequence of unobservable states that can be modeled as a Markovian process (Rabiner, 1989). The HMM assumes that the observed data have a hidden discrete-valued state sequence, and at each observed time, the observed data only depends on its hidden state. In our NOBIAS application of the HMM model, the observed data is the single-molecule displacements and the hidden state is the molecule’s distinct biophysical diffusive state.

Suppose *z*_t_ is the hidden state of the Markovian chain at time *t* and *y_t_* is the observed data at time *t*, the HMM follows the following generative process:

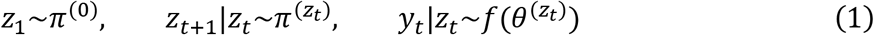

Here, *π* refers to the transition matrix of a HMM and 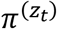 is the *z*_t_ row of the transition matrix and is the transition distribution for state *z*_t_. Given *z*_t_ and the corresponding emission parameter 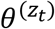, *y_t_* is independently generated from the emission function 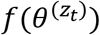. In NOBIAS, the observed data, *y_t_*, is the vector of single-step displacements, Δ***x***_t_, and the emission function is a zero-mean multivariate Gaussian distribution, and the emission parameter is the set of diffusion coefficients, 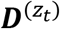:

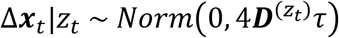

### Dirichlet process for Nonparametric Bayesian

In NOBIAS, the Dirichlet Process (DP) is used in the prior for the parameters of a mixture model with an unknown number of components. A random probability measure, *G*_0_, on a measurable space, Θ, is distributed according to a DP when (Ferguson, 1973):

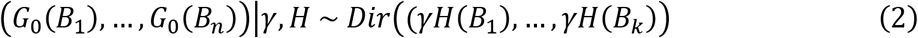

Here, *Dir* is a Dirichlet distribution, *H* is a base measurement, *γ* is a positive concentration parameter, and 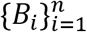 is a finite partition of Θ. In this case, we write *G*_0_~*DP*(*γ*, *H*).

From this definition follow two properties of Dirichlet processes. First, if *G*_0_~*DP*(*γ*, *H*), then *G*_0_ is atomic and can be written as:

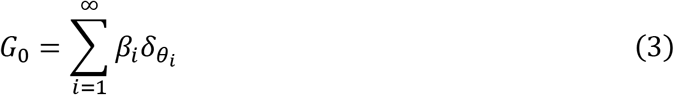

Here, *β_i_* is a weight and 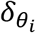 is a unit-mass measure at observation *θ_i_*|*H* ~ *H*.

Second, based on the conjugacy of the finite Dirichlet distribution, given a set of observations *θ*_1_, … , *θ*_N_ where *θ_i_*~*G*_0_, the posterior distribution for a Dirichlet process *G*_0_ is:

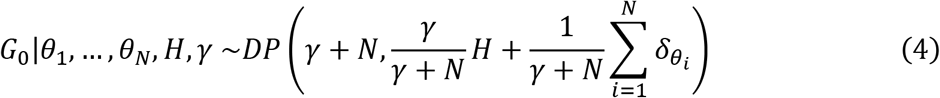

A stick-breaking process is used to construct the weight parameter *β_i_* as follows:

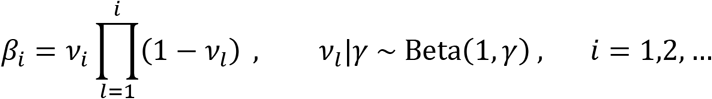

In this process, the weight *β_i_* comes from a unit stick according to a weight that is beta-distributed based on the remaining stick length after the last breaking. The weights from this construction, which is denoted *β*~GEM(*γ*), have been proven (Sethuraman, 1994) to be the weights *β_i_* of a Dirichlet process as in Eq. (3).

For each value of *θ_i_*, a random indicator variable *z*_i_ is used to denote that 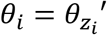, and then a predictive distribution of *z* can be written as:

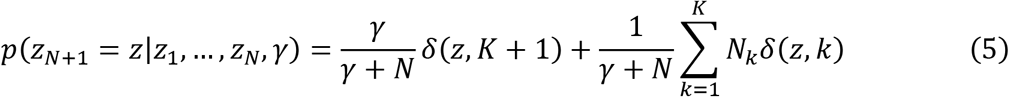

Where *K* is the current unique number of values of *z* and *N_k_* is the number of *z*_i_ that take value *k*. This predictive distribution implies that a new observation takes the value of a seen observation 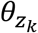 with probability proportional to *N_k_* or takes a unseen value *θ*_*K*+1_ with probability proportional to concentration parameter *γ*. When a seen observation 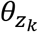 is chosen for the new observation, the indicator *z*_*N*+1_ = *k*, or if unseen value *θ*_*K*+1_ is taken, the indicator *z*_*N*+1_ = *K* + 1. This ‘the rich get richer’ property is essential for inferring a finite generated mixture model. Because the DP posterior nonparametrically converges to parameters of a mixture model for a finite mixture dataset (Ishwaran and Zarepour, 2002), the DP is an appropriate prior for the parameters of a mixture model with an unknown number of components.

### Hierarchical Dirichlet Process and Sticky Extension

In NOBIAS, the different single-molecule trajectories of multiple molecules under different biological condition and from different cells, so the groups of data are related but generated independently. Therefore, the DP is extended to a Hierarchical Dirichlet Process (HDP) (Teh et al., 2006). In the HDP, a first Dirichlet process, *G*_0_, is the base measure of a new Dirichlet process, *G_j_*:

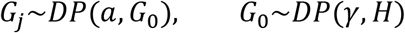

To apply a HDP as prior for a HMM model, a HDP-HMM model is generated according to:

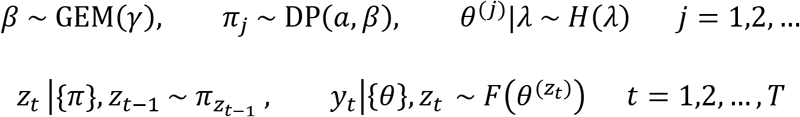

In the NOBIAS parameter setting, the observed data *y_t_* would be the single-step displacement Δ***x***_t_, the emission parameter *θ* would be the diffusion coefficient *D*, and the hyperparameter *λ* for *θ* would be the Normal-inverse-Wishart distribution (NIW) with four prior hyperparameters {*κ*, *υ*, *ν*, Δ} as stated below in the Multivariate Normal Model section.

A common issue for the HDP-HMM model is that if the algorithm artificially divides a set of observations into an alternating pattern of rapid switching between several different states, then this alternating pattern will be reinforced by the DP (Fox et al., 2008). This assignment would result in an artificial over-splitting of one state into multiple substates characterized by a high probability of transitions between the substates. Because we would not expect such rapid transitions back and forth between two distinct but similar dynamical states in the single-molecule trajectory data studied here, a sticky parameter, *κ*, is introduced which enforces self-transitions and avoids this over-splitting of states. With this new hyperparameter, the *π*_*j*_ can be sampled as:

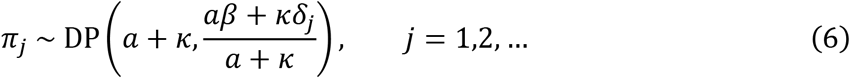

Which add a self-transition bias to the *j*^th^ components of the DP. The effects of *κ* on the results are shown in **Figure S1D**: if *κ* is too small, the over-splitting of states still occurs and if *κ* is too large, the model will underestimate the number of states.

Different Markov Chain Monte Carlo (MCMC) sampling methods such as Direct Assignment Sampling, Beam Sampling, and Blocked Sampling have been developed for the HDP-HMM model (Teh et al., 2006; Van Gael et al., 2008; Fox et al., 2007). In NOBIAS, we apply the most computationally efficient Blocked Sampling method (Fox et al., 2007), which uses a fixed-order truncation with weak-limit approximation HDP-HMM. In this approach, the DP is *L*-degree approximated as:

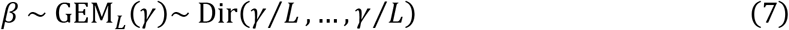

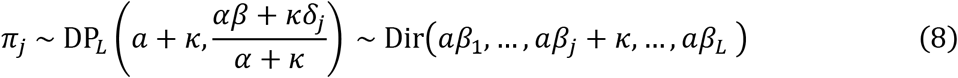

with a truncation level, *L*, that is larger than the expected total number of mixture components. Increasing *L* does not affect the posterior results, but *L* does affect the running time (**Figure S1C**). The Blocked Sampling method algorithm is detailed in the **Supplementary Note**, which describes how the state sequence is generated and how the parameters for each state are sampled.

### Multivariate Normal Model

Bayes’ rule states that the posterior distribution is proportional to the product of the prior probability and the likelihood, i.e., *P*(*θ*|*y*)~*P*(*θ*) *P*(*y*|*θ*). It is crucial to build conjugacy in order to elegantly and concisely express the posterior distribution. If we choose an appropriate prior distribution class for *P*(*θ*) given a known sampling distribution *P*(*y*|*θ*), then the posterior distribution *P*(*θ*|*y*) will have the same distribution class as the prior distribution. This choice of a prior distribution is called a conjugate prior, and this property that the posterior and prior distributions are in the same class is called conjugacy.

In NOBIAS HDP-HMM module, we assume 2D Brownian motion trajectories. In this case, the displacements follow a zero-mean 2D Gaussian and the diffusion coefficients ***D*** determine the variance, Σ, of the 2D Gaussian. Without loss of generality, the mean, *μ*, is also included in the model, *θ* = {*μ*, Σ}, and the data distribution is written as:

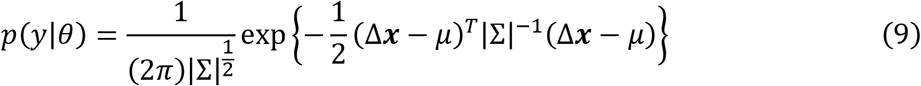

In the 2D case, the observed data, Δ***x***, is a 1 × 2 vector of the 2D displacements, *μ* is a 1 × 2 vector and Σ is the 2 × 2 covariance matrix.

As derived in reference (Gelman, 2004), the general conjugate prior model for this multivariate normal model is the prior for the mean and the variance of the step displacement follow a Normal-inverse-Wishart distribution (NIW):

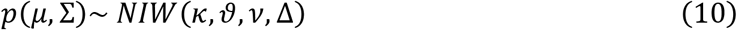

Specifically, the variance, Σ, follows an inverse-Wishart prior distribution *IW*(*ν*, Δ), and the mean, *μ*, has a conditional Normal distribution: *p*(*μ*|Σ)~ *N*(*υ*, Σ/*κ*).

The posterior updates for this normal model with NIW prior follows (Gelman, 2004):

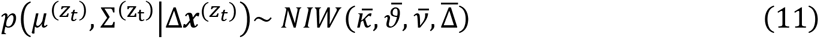

Where 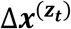 is the entire displacement dataset in state *z_t_*, and for each state *z_t_*, we update these parameters as:

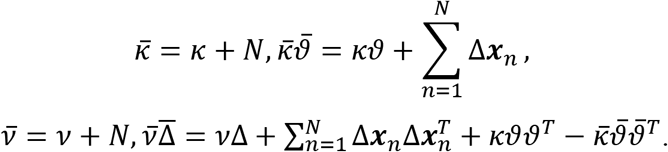

To decrease the running time, we apply the conjugate prior for the Multivariate Normal Distribution, though a non-conjugate prior is permissible. For further discussion of choice of prior see (Gelman, 2004).

### Trajectory Simulation

A state label sequence was firstly simulated with a given transition matrix through a Markov chain process. Then according the state label and the *D* of corresponding diffusive state, the 2D displacement step is generated, and cumulatively summed to get a single trajectory. Standard trajectory datasets are simulated by generate 2D Gaussian random variable where mean is 0 and variance is determined by the set diffusion coefficients with symmetry and no correlation in two directions.

Motion blur trajectory datasets are generated by simulating a state label sequence that is *T_exp_* times of the desired length with a transition matrix that self-transit enhanced *T_exp_* times. Also according to the label of this *T_exp_* times longer label sequence a true trajectories with *T_exp_* times more steps can be generated as in the standard dataset case. 2D localization error is added to the average position of every *T_exp_* steps in the true trajectory and saved to create a motion-blur trajectory with desired length. In the motion blur trajectory datasets used in this study, *T_exp_* was set to 10.

### Anomalous Diffusion

In the NOBIAS RNN module, trajectory segments of the same diffusive state (identified by the HDP-HMM module) are evaluated to classify the diffusion type for each diffusive state. In Brownian Motion, the mean squared displacement (*MSD*) is linearly proportional to the time lag, *τ*. In anomalous diffusion, *MSD* is related to *τ* according to a power law (Metzler et al., 2014):

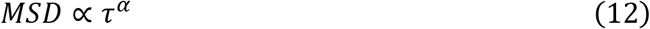

Here, *α* is the anomalous exponent. When *α* = 1, this relation describes Brownian motion; when *α* > 1, Eq. (12) describes superdiffusion; and when *α* < 1, Eq. (12) describes subdiffusion. The NOBIAS framework includes the three specific types of anomalous diffusion types that are most common in biology: Fractional Brownian motion (FBM) (Mandelbrot and Van Ness, 1968), Continuous Time Random Walk (CTRW) (Scher and Montroll, 1975), and Lévy Walk (LW) (Klafter and Zumofen, 1994).

FBM is a Gaussian process with correlated increments such that *MSD* is related to *τ* according to: 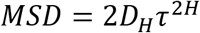 (Mandelbrot and Van Ness, 1968; Jeon and Metzler, 2010). Here, the Hurst exponent, *H*, is related to *α* in Eq. (12) by *α* = 2*H*. The *D*_*H*_ is the generalized coefficients with physical dimension 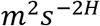. The correlation between two time points for FBM is 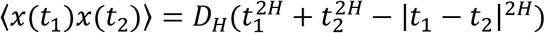. When this correlation is positive, *H* > 0.5 and the motion is superdiffusive; when the correlation is negative, *H* < 0.5 and the motion is subdiffusive.

CTRW defines a random walk family in which the particle displacement, Δ*x*, follows a wait at its current position for a random waiting time *t* that is a stochastic variable (Scher and Montroll, 1975). NOBIAS considers the case where *t* follows a power-law distribution, *ψ*(*t*) ∝ *t*^−*σ*^, and the following displacement is sampled from a zero-mean Gaussian with fixed variance. In this case, the *σ* in CTRW is related to *α* in Eq. (12), by *α* = *σ* − 1. This CTRW can only be subdiffusion, i.e., 0 < *α* ≤ 1.

LW is a special case of CTRW in which the waiting time, *t*, still follows power law, but the displacement is not Gaussian, and is instead determined by the waiting time (Klafter and Zumofen, 1994). The displacement will have a constant speed, *υ* = |Δ***x***|/*t*, and this process can only be superdiffusive with exponent 1 ≤ *α* ≤ 2.

We simulated these three types of anomalous diffusion with the open-source Python package from the recent AnDi challenge (Muñoz-Gil et al., 2020).

### Recurrent Neural Network (RNN) for NOBIAS

All segments 40 steps or greater identified in the HDP-HMM module were further analyzed by the NOBIAS Recurrent Neural Network (RNN) consisting of two long short-term memory (LSTM) layers (Hochreiter and Schmidhuber, 1997). We trained this RNN to classify trajectory segments identified to have the same diffusive state from the HDP-HMM module. We implemented this architecture, which is based on the design of the RANDI package classification task (Bo et al., 2019; Argun et al., 2021) with the MATLAB Deep Learning Toolbox™. The two LSTM layers have 100 and 50 units, respectively, and these two LSTM layers are followed by a fullyconnected layer, and the output classification layer order is given in **Figure 1**.

The input to the network is the set of 2D coordinates from the track segments; these coordinates are normalized to have zero mean and unit variance. Despite a much higher classification performance when using tracks > 50 steps long to train and validate (Argun et al., 2021; Gentili and Volpe, 2021; Muñoz-Gil et al., 2021), we trained two networks with 20-step tracks and with 40-step tracks, respectively, after considering the typical segment lengths from real biological trajectories. The training data of 750,000 trajectories were simulated with the open-source Python package from the AnDi challenge (Muñoz-Gil et al., 2020). Regression networks with similar 2 LSTM layers architecture were also trained for FBM and CTRW to estimate the anomalous exponent *α* for the experimental data. The performance of the classification network with 40-step data is shown in the confusion matrix which was made with 10000 test trajectories (**Figure S2)**. However, although the RNN module can classify CTRW and LW motion (**Figure S2**), because our HDP-HMM module assumes Brownian motion, this first module cannot predict the correct state label for these two diffusion types. We therefore test a mixture of FBM and BM motion in **Figure 3.**

### Single-Molecule Tracking in Living *Bacteroides thetaiotaomicron* Cells

*B. thetaiotaomicron* cells expressing SusG-HaloTag fusions at the native SusG promoter were grown as previously described (Karunatilaka et al., 2014). Briefly, cells were cultured overnight in 0.5% tryptone-yeast-extract-glucose medium and incubated at 37 °C under anaerobic conditions (85 % N_2_, 10 % H_2_, 5 % CO_2_) in a Coy chamber. Approximately 24 h before imaging, cells were diluted into *B. theta* minimal medium (MM) (Martens et al., 2008) containing 0.25% (wt/vol) amylopectin. On the day of the experiment, cells were diluted into fresh MM and carbohydrate and grown until reaching OD_600nm_ 0.55 – 0.60 (Tuson et al., 2018).

Before labeling, 900 μL of cells were washed twice by pelleting (6000 G, 2 min) followed by resuspension in MM. Cells were then incubated in MM supplemented with 100 nM PAJF_549_ dye (Grimm et al., 2016) for 15 min in the dark. Cells were then washed five times in MM, transferring to a new tube on every step, to remove excess dye (Lepore et al., 2019). Finally, 100 μL cells were resuspended in MM containing 0.25% (wt/vol) amylopectin for 30 min in the dark. 1.5 μL labeled cells were pipetted onto a pad of 2% agarose in MM and placed between a large and a small coverslip. The two coverslips were sealed together with epoxy (Devcon 31345 2 Ton Clear Epoxy, 25 mL) to keep the media anaerobic (Karunatilaka et al., 2014).

Cells were imaged on an Olympus IX71 inverted epifluorescence microscope with a 1.45 numerical aperture, 100× oil immersion phase-contrast objective (Olympus UPLXAPO100XOPH) and a 3.3× beam expander. Frames were collected continuously on a 512 × 512 pixel electron-multiplying charge-coupled device camera (Photometrics Evolve 512) at 50 frames/s. In this microscopy geometry, 1 camera pixel corresponds to 48.5 nm. PAJF_549_ dyes were photo-activated one at a time with a 200 – 400 ms exposure by a 406-nm laser (Coherent Cube 405-100; 0.1 μW/μm^2^) and imaged with a 561-nm laser (Coherent-Sapphire 561-50; 1 μW/μm^2^) using appropriate filters as previously described (Tuson et al., 2018).

In each movie, each cell was analyzed separately by using an appropriate mask. The collected frames were processed with SMALL-LABS (Isaacoff et al., 2019) to detect single molecules frame-by-frame and localize their position with typically ~30 nm uncertainty. Single molecules were identified as non-overlapping punctuate spots of diameter larger than 7 pixels and with pixel intensities larger than the 92^nd^ percentile intensity of the fame. The punctate spots were fit to a 2D Gaussian and true single-molecule localizations satisfied the following conditions: (1) standard deviation > 1 pixel and (2) fit error ≤ 0.06 pixel. Localizations in each cell over time were connected into trajectories using a merit value: trajectories were selected for further analysis based on their highest merit ranking.

## Results

### The NOBIAS HDP-HMM module recovers the number of diffusive states and the associated diffusion parameters

We first validated the NOBIAS HDP-HMM module with simulated single-molecule tracks, beginning from the most basic case: a mixture of Brownian motion trajectories. **Figure 2A–D** depicts the results for a mixture of two distinct diffusive states with *D*_1_ = 0.135 μm^2^/s and *D*_2_ = 1.8 μm^2^/s (**Table S1**). A sequence of state labels (1 or 2) was first simulated with a given transition matrix (probability of transitioning from state 1 to 2 or from state 2 to 1) through a Markov chain process (**Methods**). Then, according the state label and the apparent diffusion coefficient, *D*, of the corresponding diffusive state, each 2D displacement step was generated, and cumulatively summed to get a single trajectory. Similar state label sequences were simulated to generate other trajectory datasets with 4 diffusive states (**Figure 2E–G, Table S2**).

**Figure 2.**
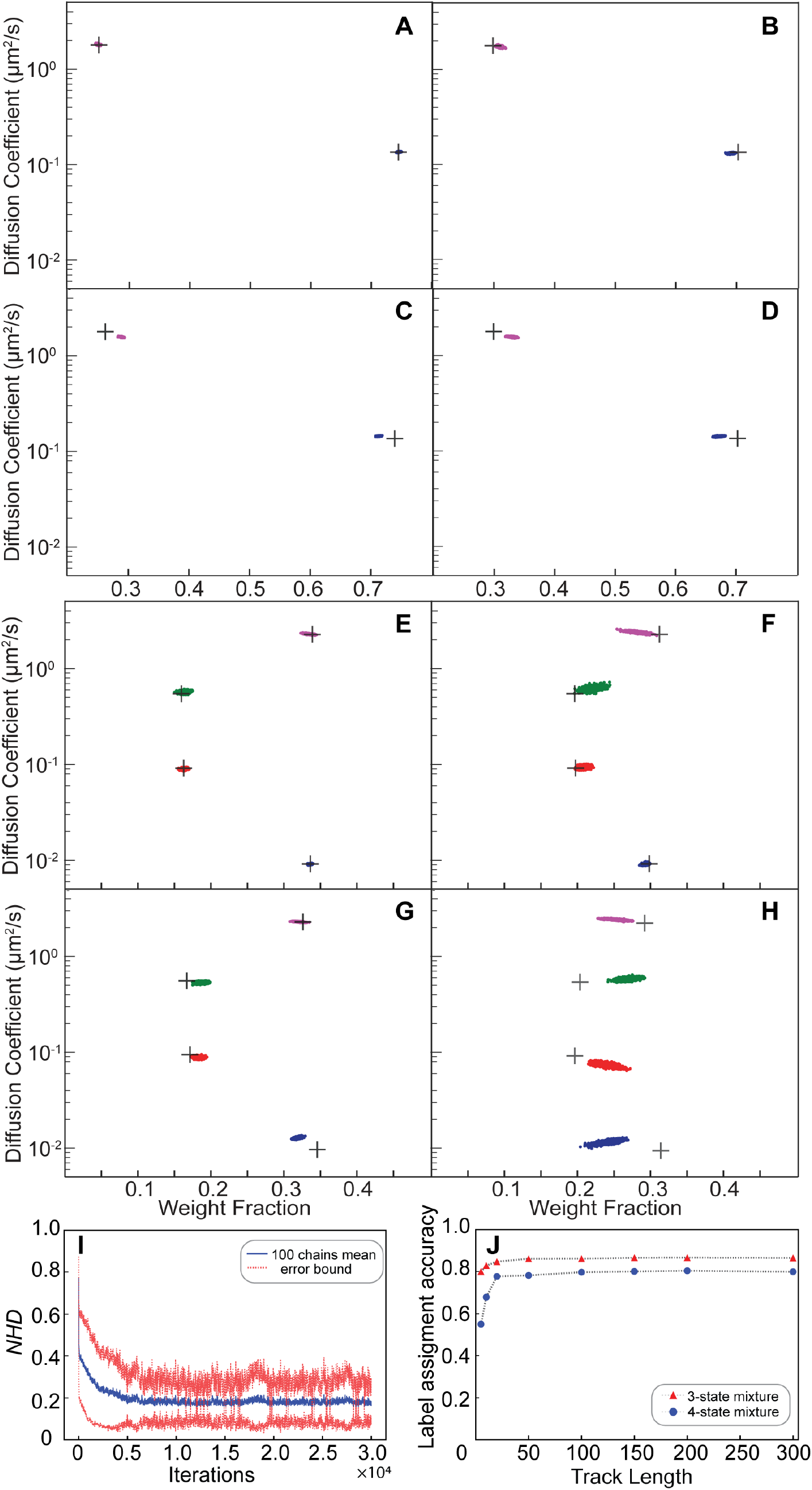
Validation of the NOBIAS HDP-HMM module with simulated trajectories. (**A-H**) The HDP-HMM module identifies distinct mobility states (colored clusters). All scatter plots include at least 500 uncorrelated samples. Each point represents the average apparent single-molecule diffusion coefficient vs. weight fraction in each distinct mobility state at each iteration of the Bayesian algorithm saved after convergence. The black crosses indicate the ground truth input for these simulated trajectories. **(A-D)** Results for two-state mixture simulated trajectories results: (**A**) Standard (no motion blur) and abundant (500 100-step trajectories) simulations, (**B**) Standard and sparse (2000 10-step trajectories) simulations, (**C**) Motion blur and abundant simulations, and (**D**) Motion blur and sparse simulations. (**E-H**) Results for four-state mixture simulated trajectories results: (**E**) Standard (no motion blur) and abundant (500 100-step trajectories) simulations, (**F**) Standard and sparse (2000 10-step trajectories) simulations, (**G**) Motion blur and abundant simulations, and (**H**) Motion blur and sparse simulations. (**I**) The normalized Hamming distance (*NHD*) decreases and converges with the number of iterations. All 100 chains use the same dataset under the settings in panel (**E**). (**J**) The final label assignment accuracy increases with the track length for three- and four-state mixture datasets. The number of trajectories decreases as the track lengths increase such that the total amount of steps is 30,000 for all track lengths.

The posterior results of the HDP-HMM module are shown in scatter plots of the inferred *D* and weight fraction from each iteration after the inferred number of states converges. **Figure 2A** shows the result for a dataset of 500 trajectories each with 100 steps. Here, the black crosses indicate the ground truth diffusion coefficient and weight fraction for each diffusive state; the posterior samples of the HDP-HMM model for the two states after convergence are distributed around the true values. Based on the posterior sample autocorrelation function (ACF) analysis (**Figure S3**), the posterior samples are thinned by saving every 10 iterations; this setting is the same for all results in this paper and was chosen by considering the effective sample sizes and the ACF analysis for all the diffusive states. The number could be updated accordingly depending on the correlation of posterior samples from output. The mean values and standard deviations for the estimation of *D* and weight fractions for the two states are listed in **Table S1**. The estimated number of unique states for this simulated dataset converges quickly over the course of iterations to the true number of states and remains mostly stable at that number (**Figure S4**). Next, we considered the less ideal case that often occurs experimentally: much shorter trajectory lengths (10 steps) and many fewer total steps (2000 10-step trajectories). We refer to the 2000 10-step trajectories as a sparse dataset and the 500 100-step trajectories are an abundant dataset. **Figure 2B** shows that the HDP-HMM model still successfully converges to the true number of states (two) for this dataset, and the posterior samples of the diffusive parameters are still distributed near the true inputs (black crosses).

We further considered the true form of collected microscope experimental data by including the localization error due to finite photon counts and noise and motion blur due to the finite image acquisition time (**Methods**). We refer these datasets ‘Motion blur dataset’ in contrast with the more ideal ‘Standard’ dataset. In the case of motion blur, the sticky parameter is increased to avoid oversampling a single diffusive state into multiple state with similar dynamics. The hyperparameter settings for this sticky HDP-HMM model are listed in **Table S3**. For both the abundant dataset (**Figure 2C**: 500 100-step trajectories) and the sparse dataset (**Figure 2D**: 2000 10-step trajectories), the true number of states (two) is recovered with our sticky HDP-HMM model, and despite these added errors, the estimated parameters deviate only slightly from the true inputs (black crosses).

We extended our simulations of standard and motion blur Brownian motion track mixtures to a more complicated 4-state scenarios for abundant (500 100-step trajectories) and sparse (2000 10-step trajectories) datasets (**Figure 2 E–H**). Even with 4 diffusive states, the performance of the HDP-HMM module is still excellent for the standard mixture (**Figure 2E–F**). For the 4-state mixture simulation that includes localization error and motion blur, the HDP-HMM still successfully recovers the true number of states, and the parameters for the four distinct states are still estimated well, though the posterior samples have increased variance and deviation from the true value (**Figure 2G–H**). The statistics of the posterior samples for estimated parameters of the 4-state simulation result are listed in **Table S2**, and the transition matrices for all the simulations in **Figure 2** are shown in **Table S1-S2**.

The NOBIAS HDP-HMM module also assigns diffusive state labels to each single-molecule step within the trajectories dataset; we call this the state sequence for each track. We quantified the performance of the state sequence assignment relative to the ground truth simulated state sequence with the Hamming distance: the Hamming distance between two 1D sequences with equal length is the number of points where the components are different (Hamming, 1950). The resulting distances were normalized to the total length to demonstrate the Normalized Hamming Distance (*NHD*) convergence over iterations (**Figure 2I**). The *NHD* decreases with increasing iteration number and converges to approximately 0.18. This final converged *NHD* depends on the dataset size, the true transition matrix, and how separable the diffusive state are from one another.

The true number of diffusive states can be recovered for datasets of both abundant and sparse tracks, but the HDP-HMM module performance depends strongly on the length of the individual tracks. Using the overall state sequence assignment accuracy (1 - *NHD*) as a performance evaluator for datasets with the same total amount of steps (30000), we found that the assignment accuracy is considerably worse for tracks shorter than 20 steps and almost linearly increases with the track length till asymptotes for longer tracks (> 20 steps; **Figure 2J**). This trend is shared for a 3-state and 4-state dataset, but the overall accuracy for 3-state dataset is higher than 4-state one for all the track length.

### The NOBIAS RNN module predicts the diffusion type for each diffusive state

To analyze anomalous diffusion in an SPT dataset, NOBIAS includes a second module: we built an RNN to classify the type of motion (Brownian motion (BM), Fractional Brownian motion (FBM), Continuous Time Random Walk (CTRW), or Lévy Walk (LW)) corresponding to the track segments within each diffusive state identified by HDP-HMM module. The RNN consists of two LSTM layers, a fullyconnected layer, and data input/output layer (**Methods**). Although the HDP-HMM module is based on BM, for some anomalous diffusion types, for example FBM, if the dynamics level for each state is distinct, the HDP-HMM module still performs well.

We simulated a mixture of BM and FBM with distinct apparent diffusion coefficients for the two states (*D*_1_ = 0.045 μm2/s and *D*_2_ = 0.90 μm2/s) to validate the performance of NOBIAS on mixtures of different diffusion types. **Figure 3A** shows the HDP-HMM posterior results for this 2-state BM-FBM mixture (500 100-step trajectories) where the FBM state is anomalous subdiffusion with *α* = 0.5 (Eq. (12)) and with lower diffusion coefficient. Then, based on the state sequence labels from the HDP-HMM module, we generated track segments for the two diffusive states and put them into the trained NOBIAS RNN network to predict the diffusion types. NOBIAS RNN successfully predicts the diffusion types for both states (**Figure 3B, Table S4**).

**Figure 3.**
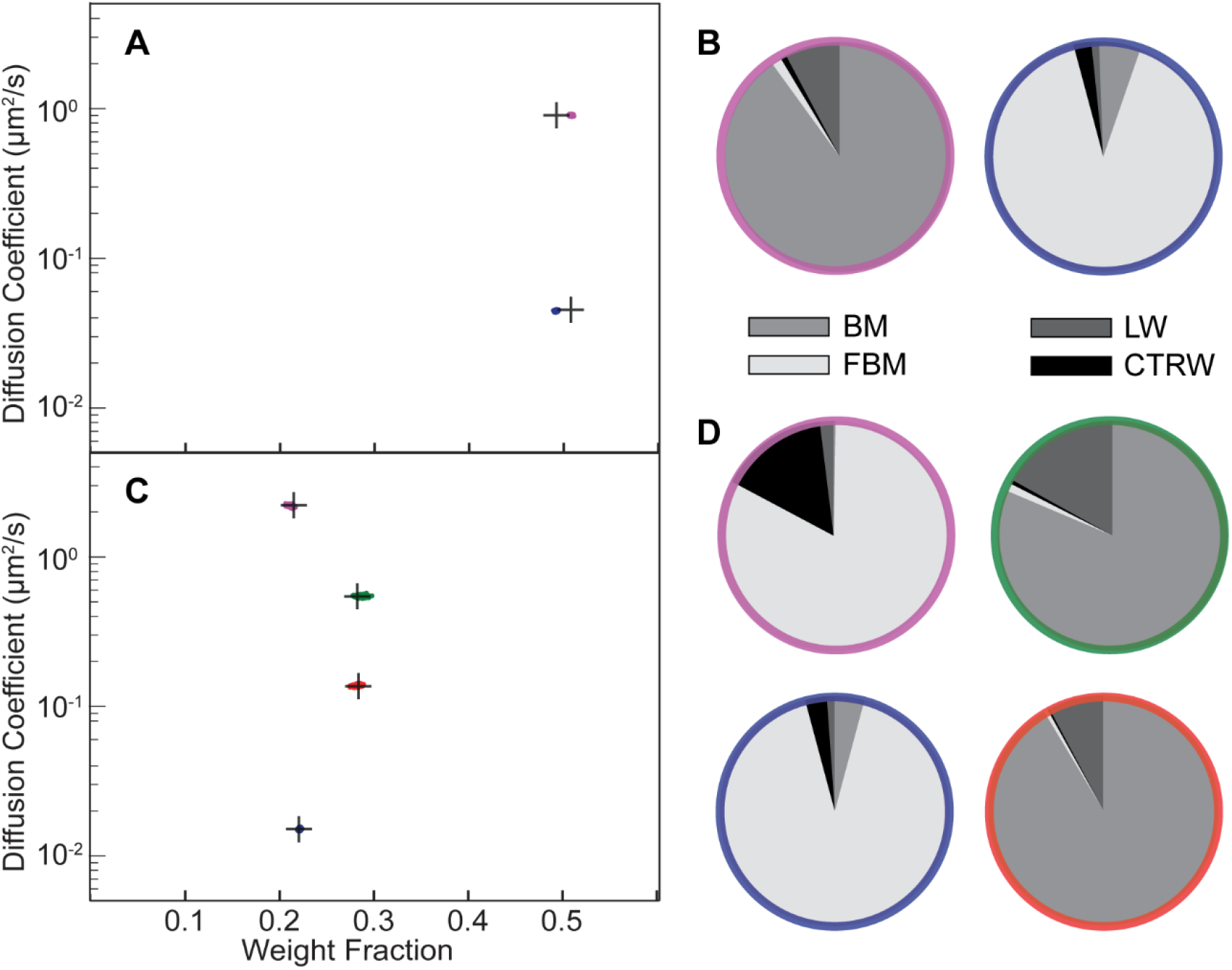
Validation of the NOBIAS-RNN module with simulated trajectories containing mixtures of different diffusion types. (**A, C**) The HDP-HMM module identifies distinct mobility states (colored clusters). Each point represents the average apparent single-molecule diffusion coefficient, *D*, vs. weight fraction in each distinct mobility state at each iteration of the Bayesian algorithm saved after convergence. The black crosses indicate the ground truth input for these simulated trajectories. (**A**) Two-state mixture comprising a subdiffusive Fractional Brownian Motion (FBM) state with lower *D* and a Brownian Motion (BM) state with higher *D*. (**B**) The NOBIAS-RNN determines the probability that the diffusion type for each diffusive state in (**A**) is classified as BM, FBM, Continuous Time Random Walk (CTRW), or Lévy Walk (LW). The final probability for each diffusive state is the average of the classification probability of its track segments weighted by the segment length. The color of each pie chart indicates the diffusive state corresponding to the color in (**A**). (**C**) Four-state mixture comprising a subdiffusive FBM state, two BM states, and a superdiffusive FBM state with *D* in ascending order. (**D**) Diffusion type classification probability pie chart for each diffusive state in (**C**). The final probability for each diffusive state is the average of the classification probability of its track segments weighted by the segment length and the color of each pie chart indicates the diffusive state corresponding to the corresponding color in (**C**).

We further simulated a 4-state mixture (500 100-step trajectories) corresponding to subdiffusive FBM, BM, BM, and superdiffusive FBM (in order of increasing *D*). The HDP-HMM module still successfully recovers the 4 states and make excellent estimations for *D* and weight fraction for each state (**Figure 3C**). The NOBIAS RNN module also predicts the true diffusion type for the segments from each of the four states (**Figure 3D, Table S4**). Note that all track segments are normalized before being put into the RNN to avoid dynamics information bias in the diffusion type prediction (**Methods**). One limitation for this RNN classification analysis methodology is that only track segments with at least certain length (20 or 40 in our analysis depending on the trained network) could be classified with high accuracy; it is very challenging to use very short track segments to identify these modes of diffusion. Therefore, when the overall trajectory length is short (~10 steps), the network classification module might not be usable. Another limitation of the HDP-HMM module is that the current implementation is based on BM displacement distributions, thus it would fail for anomalous diffusion types like LW, which does not have a similar Gaussian distribution of displacements.

### Performance of NOBIAS on experimental data for the diffusion of SusG-HaloTag in *Bacteroides thetaiotaomicron* cells

After validating the performance of the two NOBIAS modules on simulated data, we applied this framework to experimental single-molecule trajectories. The SusG amylase recognizes and binds starch on the surface of *B. thetaiotaomicron* cells to enable starch catabolism (Koropatkin and Smith, 2010). We measured the motion of 7897 trajectories (minimum length of 6 and average length of 64) of single SusG molecules in 226 *B. thetaiotaomicron* cells based on imaging photoactivatable fluorescently labeled SusG-HaloTag fusions (**Methods**).

We analyzed this data with NOBIAS to infer the number of diffusive states and to estimate the diffusion coefficient, weight fraction, and type of motion for each state as was done for the simulated data (**Figure 2 and 3**). Additionally, NOBIAS analyzes 2D trajectories with a 2D Gaussian function and can therefore infer the diffusion coefficients for the *x* and *y* directions separately and estimate the potential correlation between the two directions. Though the simulations used symmetric tracks in an unbound domain, the experiments measure motion on the surface of cells with a long axis and a short axis, which may create an asymmetry in the diffusion. We rotated the cell orientations to orient the long axis in the *x* direction without rescaling (**Figure 4A**). We analyzed this rotated dataset with NOBIAS and found that it converged to a 3-state model, with a very small (1.8%) fast state fraction (**Figure 4B**). Interestingly, we found that the *D*_x_ and *D*_y_ values were similar for each of the two slower states (**Table S5**), while they were significantly different for the fastest state (*D*_x_ = 0.68 μm2/s vs. *D*_y_ = 0.45 μm2/s). This asymmetry for the fast state indicates that it corresponds to free diffusion that is constrained by the cell shape (and therefore is more constrained in the short-axis y direction), while the symmetry for the two slower states implies molecules that only diffuse regionally and are not affected by the cell shape. Compared with previous SPT analysis methods, NOBIAS provides a two-dimensional analysis of the dynamics of experimental single-molecule trajectories.

**Figure 4.**
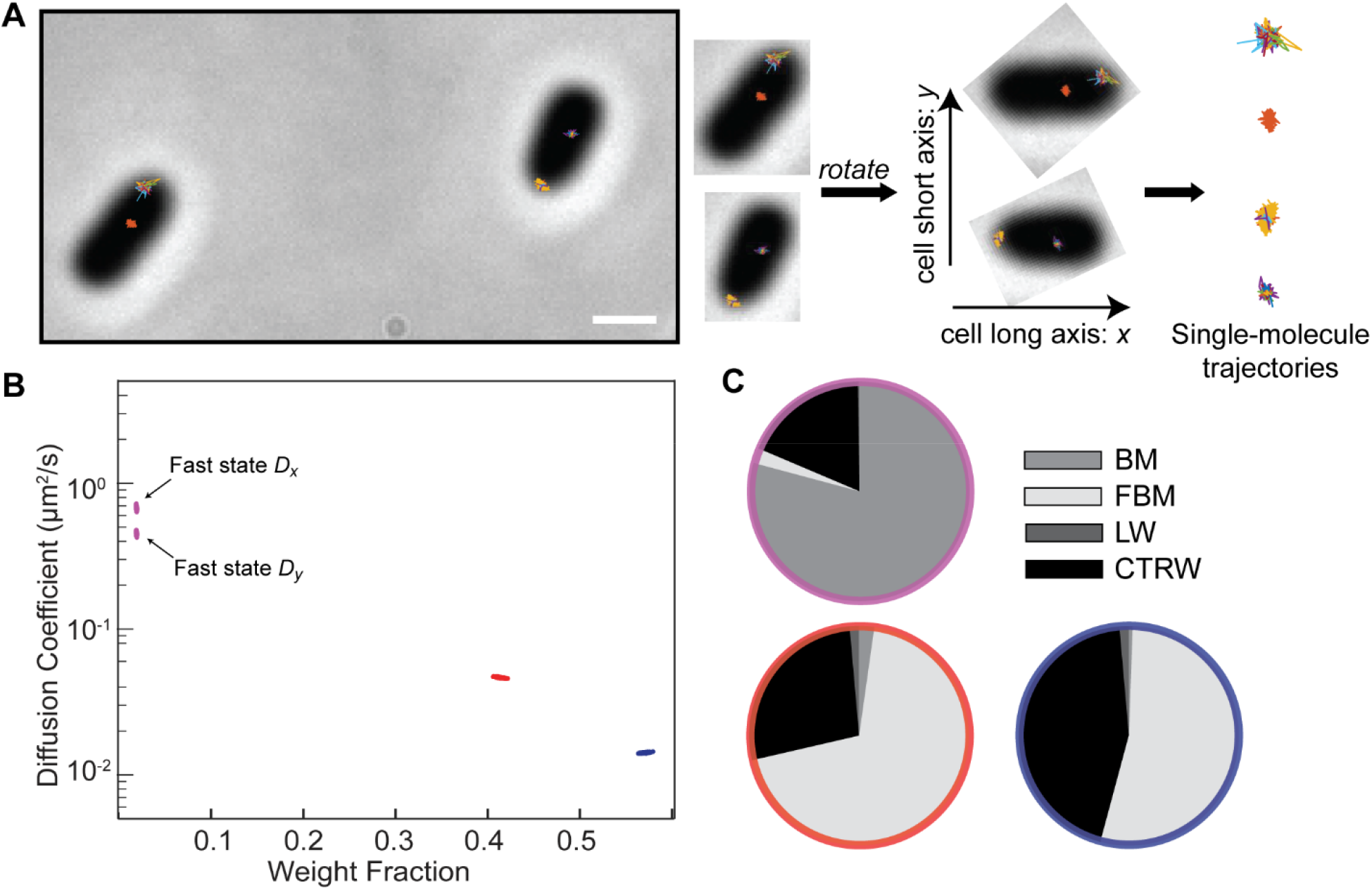
Application of NOBIAS to single-molecule trajectories of the SusG protein in living *Bacteroides thetaiotaomicron* cells. (**A**) Single-molecule trajectories of SusG-HaloTag overlaid on the phase-contrast image of the corresponding *B. thetaiotaomicron* cells, scale bar: 1 *μ*m. The long axis of the phase mask for each cell was detected and a rotation transform was applied to all the trajectories in each cell such that the *x*-axis is the cell long axis for all cells. (**B**) The NOBIAS HDP-HMM module identifies three diffusive states for SusG (colored clusters). Each point represents the average apparent single-molecule diffusion coefficient vs. weight fraction in each distinct mobility state at each iteration of the Bayesian algorithm saved after convergence. The blue and red points clusters average the x- and y- diffusion coefficients as they are symmetric (Table S4); the asymmetric fast state (purple) shows a different **D*_x_* and **D*_y_*. (**C**) The NOBIAS-RNN determines the probability that the diffusion type for each diffusive state in (**B**) is classified as Brownian Motion (BM), Fractional Brownian Motion (FBM), Continuous Time Random Walk (CTRW), or Lévy Walk (LW). The color of each pie chart indicates the diffusive state corresponding to the color in (**B**). The fast state (purple) is predicted with high probability to be BM; the two slower states (red and blue) are predicted to be FBM or CTRW.

We separated the track segments by the state sequence label from the HDP-HMM module and placed each group into the RNN classification module. The fastest state was predicted with high probability (80%) to be Brownian motion (**Figure 4C, Table S4**), consistent with the asymmetry between *D_x_* and *D_y_* that was attributed to free diffusion (**Figure 4B**). The two slower states were predicted to be either FBM or CTRW. We used a RNN regression network (**Methods**) to estimate the anomalous exponent *α* for the track segments of the two slower states and both were found to be subdiffusion (*α*_1_ = 0.38, *α*_2_ = 0.46), consistent with the symmetry between *D_x_* and *D_y_* found (**Table S5**). This finding of subdiffusion is also consistent with the role of SusG in starch catabolism: we have previously found that SusG motion slows in the presence of its amylopectin substrate, as well as when it transiently associates other outer-membrane proteins, indicating starch-mediated Sus complex formation (Karunatilaka et al., 2014).

## Discussion

Single-molecule tracking measures dynamics in biological systems at high spatial and temporal resolution, but how to make the best use of these tracking data for a broad set of experimental conditions remains an analysis challenge in the field (Shen et al., 2017; Elf and Barkefors, 2019). Here, we have introduced NOBIAS to quantify single-molecule dynamics and to associate these biophysical measurements with the underlying biochemical function and biological processes. NOBIAS handles complicated live-cell SPT datasets for which: (1) the number of diffusive states is unknown, (2) mixtures of different diffusive populations may exist, even within single trajectories, (3) symmetry cannot be assumed between the *x* and *y* directions, and (4) anomalous diffusion is possible. These features are enabled based on applying Nonparametric Bayesian statistics (Teh et al., 2006; Fox et al., 2008; Johnson and Willsky, 2013) to SPT datasets that have the same means but different variance with a HDP-HMM module that has a 2D Gaussian as the emission function and then by further investigating the anomalous diffusion types in the RNN module of NOBIAS.

Compared with previous applications of nonparametric Bayesian statistics in this field (Persson et al., 2013; Karslake et al., 2020; Heckert et al., 2021), the NOBIAS HDP-HMM module is more robust and has high computational efficiency (**Table S6**). NOBIAS and SMAUG both consider motion blur effects and their estimation of *D* for each state is closer to the ground truth then other methods. As Bayesian method with similar principle NOBIAS is almost 10 times faster than SMAUG. This HDP-HMM module also provides a multivariate output to quantify and correlate dynamics in multiple directions instead of assuming symmetry (**Table S7**). We observed that for asymmetric simulated trajectories, vbSPT overestimates the true number of states, and SMAUG can only provide the average *D* of for each diffusive state while NOBIAS provides the respective diffusion coefficients in two directions. The high accuracy of step state sequence prediction also enables the classification of anomalous diffusion type in the NOBIAS RNN module. We also applied SMAUG and vbSPT on the experimental dataset (**Table S8**): SMAUG ran slow on large datasets and suggested four diffusive state, while vbSPT suggested the best model to be 10 diffusive state which is hard to explain their corresponding biological meanings.

A further advantage of NOBIAS lies in its ability to treat sets of relatively short trajectories (10-step trajectories in the simulated data of **Figures 2 and 3** and minimal 6-step trajectories in the experimental data of **Figure 4**). The recent AnDi (Anomalous Diffusion) Challenge (Muñoz-Gil et al., 2021) demonstrated that Deep Learning and Neural Network methods are currently the most powerful tools to study anomalous diffusion (Argun et al., 2021; Gentili and Volpe, 2021). However, in this challenge, the target dataset was an ideal collection of simulated anomalous diffusion trajectories with 100-1000 steps, and only the simple case of one state transition in the middle part of a track was considered. There are also probability-based models that consider confinement and anomalous diffusion (Robson et al., 2013) and Bayesian methods that directly predict the diffusion type (Thapa et al., 2018; Cherstvy et al., 2019), but these analyses, like the neural network-based methods, are used for very long trajectories or assume a single diffusive state in each track. To apply a deep learning-based diffusion type classifier to realistic simulated trajectories and real experimental trajectories, NOBIAS segments the raw trajectories into collections of track segments that belong to the same diffusive state (as identified by the HDP-HMM module) and then predicts the diffusion type of the long segments in the RNN module. Since different biophysical diffusive states correspond to different biochemical functions which will exhibit different diffusion types due to interactions like confinement, binding, directional motion, NOBIAS enables a thorough investigation of these biochemical roles by revealing the diffusion coefficients, the transition probabilities between states, and the anomalous diffusion behaviors. Ultimately, NOBIAS will enable investigators to extract a complete information set from SPT data and to understand the role of each tracked molecule, even in the living cell.

Despite these strengths, NOBIAS has several limitations. Firstly, as an HMM-based method, NOBIAS is limited by the length of each track. Under the extreme case where only very short trajectories (~2-5 steps) are available, the HDP-HMM module may suggest a number of states and posterior results with extremely high uncertainty; probability-based models (Rowland and Biteen, 2017; Hansen et al., 2018) or the histogram-based Bayesian method DPMM (Heckert et al., 2021) should be applied for these short trajectories. The track length also limits the RNN module, as the trained network need tracks with at least 20 steps for good classification performance because some anomalous diffusion types are characterized by memory of previous steps (Metzler et al., 2014). Therefore the application of the RNN module is limited for short experimental tracks. Secondly, NOBIAS performs the diffusive state estimation based on apparent diffusion coefficient in the HDP-HMM module and then carries out the anomalous diffusion classification in the RNN module. NOBIAS therefore assumes that each biochemical state has a unique average apparent diffusion coefficient. Although the RNN module can classify the diffusion types of two different diffusive states with the same diffusion coefficient, the HDP-HMM module would fail to separate these processes. Furthermore, for some diffusion types like LW, the trajectory displacements may exhibit different types of dynamics even though the trajectories are generated from one process. Finally, even for Brownian trajectories, a single biochemical state might not be represented by a single diffusion coefficient value. Thus, the actual number of biochemical states may not be equal to the number of diffusive states. Future development of NOBIAS could use spatial filtering to distinguish between these similar biochemical states.

NOBIAS provides a pioneering and compatible framework for the analysis of dynamical mixtures that also classifies the anomalous diffusion types. Future development of NOBIAS could include more types of diffusion and could integrate the anomalous distributions directly into the Bayesian framework for more accurate prediction of the stepwise state labels and the diffusion types. Furthermore, extra experimental corrections corresponding to the specific microscope setting (Berglund, 2010; Lindén et al., 2017; Hansen et al., 2018) could also help adapt NOBIAS more broadly to different types of SPT datasets. Overall, NOBIAS has provided a powerful framework to analyze of SPT dataset with unknown number of diffusive states and potential asymmetric diffusion, and to access the anomalous diffusion type for each diffusive state. The combination of nonparametric Bayesian statistics and Deep learning enables NOBIAS to fully extract the rich dynamics information from the SPT dataset.

## Supplementary Materials

Supplementary Figures S1 – S4, Supplementary Tables S1 – S8, and a Supplementary Note are provided; the raw data and outputs investigated in this manuscript are also provided. Open-source Matlab code for implementing NOBIAS (GNU General Public License) and some test datasets are provided at https://github.com/BiteenMatlab/NOBIAS; further development and expansion of the code post-publication will be hosted at that website as well.

## Author Contributions

Z.C. and J.S.B. conceived of the idea. Z.C. developed the theory, implemented the algorithm, performed simulations, and analyzed simulated and experimental data. L.G. carried out the experiments. Z.C. and J.S.B. wrote the manuscript with input from all authors.

## Acknowledgements

This work was supported by National Institutes of Health grant R21-GM128022 and National Science Foundation grant EF-1921677 to JSB. Thanks to Christopher Azaldegui and Guoming Gao for helpful discussions.

