## Supplemental Data 1 for "NOBIAS: Analyzing anomalous diffusion in single-molecule tracks with nonparametric Bayesian inference"

### Supplementary Material for
### NOBIAS: Analyzing anomalous diffusion in single-molecule tracks with nonparametric Bayesian inference

Ziyuan Chen[1], Laurent Geffroy[2], Julie Biteen[1,2,*]

University of Michigan, Departments of [1]Biophysics and [2]Chemistry, Ann Arbor, MI 48109

*

**Contents:**

- SI Tables S1 – S8
- SI Figures S1 – S4
- SI Note: Blocked Sampler for the NOBIAS HDP-HMM module

**Table S1**. Two-state mixture results for simulations of the standard abundant, standard sparse, motion blur abundant, motion blur sparse, and a mixture of Brownian and subdiffusive fractional Brownian motion models. Diffusion coefficients (in $\mu m^2/s$) and weight fractions (in %) are given for the ground truth inputs and the NOBIAS HDP-HMM module outputs for each of the 2 states. These data correspond respectively to the main text figures as indicated. Errors represent standard deviation.

| | **State 1 ($\mu m^2/s$; %)** | **State 2 ($\mu m^2/s$; %)** | **Main Text Figure** |
|---|---|---|---|
| Standard abundant | 0.135<br>74.75 | 1.8<br>25.25 | 2A |
| NOBIAS | $0.136 \pm 0.001$<br>$74.82 \pm 0.14$ | $1.824 \pm 0.019$<br>$25.18 \pm 0.14$ | 2A |
| Standard sparse | 0.135<br>70.23 | 1.8<br>29.78 | 2B |
| NOBIAS | $0.132 \pm 0.001$<br>$69.12 \pm 0.26$ | $1.754 \pm 0.026$<br>$30.88 \pm 0.26$ | 2B |
| Motion blur abundant | 0.135<br>74.32 | 1.8<br>25.68 | 2C |
| NOBIAS | $0.143 \pm 0.001$<br>$71.35 \pm 0.18$ | $1.579 \pm 0.010$<br>$28.65 \pm 0.18$ | 2C |
| Motion blur sparse | 0.135<br>70.24 | 1.8<br>29.76 | 2D |
| NOBIAS | $0.142 \pm 0.001$<br>$67.20 \pm 0.32$ | $1.580 \pm 0.016$<br>$32.80 \pm 0.32$ | 2D |
| Mixture of BM and FBM | 0.045<br>50.76 | 0.90<br>49.24 | 3A |
| NOBIAS | $0.044 \pm 0.0003$<br>$49.20 \pm 0.08$ | $0.901 \pm 0.006$<br>$50.80 \pm 0.08$ | 3A |

**Table S2**. Four-state mixture results for simulations of the standard abundant, standard sparse, motion blur abundant, motion blur sparse, and a mixture of Brownian, subdiffusive fractional Brownian, and superdiffusive fractional Brownian motion models. Diffusion coefficients (in $\mu m^2/s$) and weight fractions (in %) are given for the ground truth inputs and the NOBIAS HDP-HMM module outputs for each of the 4 states. These data correspond respectively to the main text figures as indicated. Errors represent the standard deviation.

| | **State 1 ($\mu m^2/s$; %)** | **State 2 ($\mu m^2/s$; %)** | **State 3 ($\mu m^2/s$; %)** | **State 4 ($\mu m^2/s$; %)** | **Main Text Figure** |
|---|---|---|---|---|---|
| Standard abundant | 0.009<br>33.70 | 0.09<br>16.32 | 0.54<br>16.01 | 2.25<br>33.97 | 2E |
| NOBIAS | $0.009 \pm 0.0001$<br>$33.68 \pm 0.11$ | $0.089 \pm 0.002$<br>$16.37 \pm 0.20$ | $0.561 \pm 0.012$<br>$16.44 \pm 0.36$ | $2.275 \pm 0.022$<br>$33.51 \pm 0.33$ | 2E |
| Standard sparse | 0.009<br>29.74 | 0.09<br>19.61 | 0.54<br>19.51 | 2.25<br>31.14 | 2F |
| NOBIAS | $0.009 \pm 0.0001$<br>$29.18 \pm 0.22$ | $0.092 \pm 0.003$<br>$20.79 \pm 0.41$ | $0.610 \pm 0.026$<br>$21.92 \pm 0.72$ | $2.371 \pm 0.048$<br>$28.11 \pm 0.74$ | 2F |
| Motion blur abundant | 0.009<br>34.74 | 0.09<br>16.64 | 0.54<br>16.16 | 2.25<br>32.46 | 2G |
| NOBIAS | $0.012 \pm 0.0002$<br>$31.74 \pm 0.29$ | $0.084 \pm 0.002$<br>$18.04 \pm 0.35$ | $0.518 \pm 0.009$<br>$18.32 \pm 0.41$ | $2.263 \pm 0.015$<br>$31.89 \pm 0.36$ | 2G |
| Motion blur sparse | 0.009<br>31.32 | 0.09<br>19.44 | 0.54<br>20.15 | 2.25<br>29.10 | 2H |
| NOBIAS | $0.011 \pm 0.0001$<br>$24.32 \pm 1.08$ | $0.073 \pm 0.003$<br>$23.72 \pm 0.99$ | $0.578 \pm 0.016$<br>$26.7 \pm 0.80$ | $2.441 \pm 0.036$<br>$25.25 \pm 0.80$ | 2H |
| Mixture of BM and FBM | 0.015<br>22.01 | 0.135<br>28.31 | 0.54<br>28.20 | 2.21<br>21.49 | 3C |
| NOBIAS | $0.015 \pm 0.0002$<br>$22.09 \pm 0.07$ | $0.136 \pm 0.001$<br>$28.15 \pm 0.20$ | $0.541 \pm 0.006$<br>$28.66 \pm 0.26$ | $2.191 \pm 0.024$<br>$21.10 \pm 0.17$ | 3C |

**Table S3.** NOBIAS HDP-HMM module hyperparameter settings for the simulations and experimental data in main text Figures 2 – 4.

| **Hyperparameter** | **Standard** | **Motion blur abundant** | **Motion blur sparse** | **Experimental** |
|---|---|---|---|---|
| $\gamma$ | 0.1 | 0.1 | 0.1 | 0.1 |
| $a$ | 1 | 1 | 1 | 1 |
| $\kappa$ | 5 | 100 | 10 | 100 |

**Table S4.** NOBIAS RNN module diffusion type classification probabilities for Brownian motion (BM), Fractional Brownian motion (FBM), Continuous Time Random Walk (CTRW), and Lévy Walk (LW). These data correspond respectively to the main text figures as indicated.

| | BM (%) | FBM (%) | CTRW (%) | LW (%) | Main Text Figure |
|---|---|---|---|---|---|
| Mixture of BM and FBM | 6.04 | 90.35 | 2.71 | 0.90 | 3B |
| | 90.06 | 1.57 | 0.66 | 7.71 | |
| Mixture of Four Anomalous Diffusion Types | 4.41 | 90.89 | 3.63 | 1.07 | 3D |
| | 91.44 | 0.59 | 0.23 | 7.73 | |
| | 81.60 | 0.92 | 0.75 | 16.73 | |
| | 0.33 | 82.59 | 15.26 | 1.82 | |
| SusG-HT Experimental Data | 0.75 | 53.67 | 44.27 | 1.31 | 4C |
| | 2.36 | 69.10 | 27.27 | 1.28 | |
| | 79.17 | 2.39 | 18.30 | 0.13 | |

**Table S5.** NOBIAS HDP-HMM module results for analysis of the experimental measurements of SusG-HT diffusion in the *Bacteroides thetaiotaomicron* outer membrane corresponding to main text Figure 4B. The *x*-axis and *y*-axis diffusion coefficients ($D_x$ and $D_y$, respectively) are evaluated separately in NOBIAS.

| | State 1 | State 2 | State 3 | Transition Matrix |
|---|---|---|---|---|
| $D_x$ ($\mu m^2/s$) | $0.013 \pm 0.0001$ | $0.043 \pm 0.0003$ | $0.675 \pm 0.014$ | $\begin{bmatrix} 0.933 & 0.067 & 0 \\ 0.098 & 0.897 & 0.005 \\ 0.006 & 0.148 & 0.846 \end{bmatrix}$ |
| $D_y$ ($\mu m^2/s$) | $0.015 \pm 0.0001$ | $0.049 \pm 0.0003$ | $0.450 \pm 0.009$ | |
| Weight (%) | $56.94 \pm 0.24$ | $41.29 \pm 0.23$ | $1.77 \pm 0.02$ | |

**Table S6.** Comparison of results for analyzing simulated Brownian motion data with symmetric diffusion coefficient inputs in NOBIAS and three established nonparametric Bayesian statistics algorithms. The truncation level of NOBIAS, the max state number of vbSPT, and the starting number of states in SMUAG were all set to 10.

| | **Number of States** | **Processing Time (s)** | **State 1 ($\mu m^2/s$) (%)** | **State 2 ($\mu m^2/s$) (%)** | **State 3 ($\mu m^2/s$) (%)** | **Transition Matrix** | **Reference** |
|---|---|---|---|---|---|---|---|
| Ground Truth Input | 3 | - | 0.015<br>37.36 | 0.15<br>23.94 | 1.5<br>38.70 | $\begin{bmatrix} 0.9 & 0.05 & 0.05 \\ 0.1 & 0.8 & 0.1 \\ 0.05 & 0.05 & 0.9 \end{bmatrix}$ | - |
| NOBIAS | 3 | 925 | 0.0154<br>33.53 | 0.1500<br>27.90 | 1.4878<br>38.56 | $\begin{bmatrix} 0.90 & 0.07 & 0.03 \\ 0.08 & 0.82 & 0.10 \\ 0.03 & 0.06 & 0.91 \end{bmatrix}$ | current |
| vbSPT | 3 | 364 | 0.0119<br>35.18 | 0.1076<br>25.90 | 0.9891<br>38.92 | $\begin{bmatrix} 0.91 & 0.06 & 0.03 \\ 0.09 & 0.82 & 0.09 \\ 0.03 & 0.05 & 0.92 \end{bmatrix}$ | Persson 2013 |
| SMAUG | 3 | 11138 | 0.0161<br>36.50 | 0.1519<br>26.53 | 1.4776<br>38.91 | $\begin{bmatrix} 0.89 & 0.08 & 0.04 \\ 0.11 & 0.78 & 0.11 \\ 0.04 & 0.07 & 0.89 \end{bmatrix}$ | Karslake 2020 |
| Spot-On | 3* | 33.8 | 0.011<br>27.1 | 0.076<br>33.7 | 0.883<br>39.2 | NA | Hansen 2018 |

*The number of states was fixed at 3 for Spot-On.

**Table S7.** Comparison of results for analyzing simulated Brownian motion data with asymmetric diffusion coefficient inputs in NOBIAS and three established nonparametric Bayesian statistics algorithms. The truncation level of NOBIAS, the max state number of vbSPT, and the starting number of states in SMUAG were all set to 10.

| | **Number of States** | **State 1 ($\mu m^2/s$) (%)** | **State 2 ($\mu m^2/s$) (%)** | **State 3 ($\mu m^2/s$) (%)** | **Transition Matrix** | **Reference** |
|---|---|---|---|---|---|---|
| Ground Truth Input | 3 | $D_x$: 0.021<br>$D_y$: 0.009<br>36.65 | $D_x$: 0.21<br>$D_y$: 0.09<br>24.49 | $D_x$: 2.1<br>$D_y$: 0.9<br>38.86 | $\begin{bmatrix} 0.9 & 0.05 & 0.05 \\ 0.1 & 0.8 & 0.1 \\ 0.05 & 0.05 & 0.9 \end{bmatrix}$ | - |
| NOBIAS | 3 | $D_x$: 0.0216<br>$D_y$: 0.0094<br>32.82 | $D_x$: 0.2025<br>$D_y$: 0.0918<br>28.11 | $D_x$: 2.0657<br>$D_y$: 0.8868<br>39.06 | $\begin{bmatrix} 0.90 & 0.07 & 0.03 \\ 0.08 & 0.82 & 0.10 \\ 0.03 & 0.06 & 0.91 \end{bmatrix}$ | current |
| vbSPT | 4 | 0.0115<br>34.64 | 0.1010<br>24.31 | 0.5191<br>9.27<br>† | $\begin{bmatrix} 0.91 & 0.05 & 0.03 & 0.01 \\ 0.09 & 0.80 & 0.07 & 0.04 \\ 0.07 & 0.03 & 0.56 & 0.34 \\ 0.01 & 0.06 & 0.16 & 0.77 \end{bmatrix}$ | Persson 2013 |
| SMAUG | 3 | 0.0153<br>34.04 | 0.1493<br>27.32 | 1.4996<br>38.64 | $\begin{bmatrix} 0.878 & 0.086 & 0.036 \\ 0.11 & 0.75 & 0.14 \\ 0.035 & 0.09 & 0.875 \end{bmatrix}$ | Karslake 2020 |
| Spot-On | 3* | 0.011<br>27.4 | 0.073<br>33.2 | 0.83<br>39.5 | NA | Hansen 2018 |

* The number of states was fixed at 3 for Spot-On.

† The vbSPT analysis indicated a 4th state with $D$ = 1.2249 $\mu m^2/s$ and weight fraction = 25.38%.

**Table S8.** Comparison of results for analyzing experimental data in NOBIAS and two established nonparametric Bayesian statistics algorithms. The truncation level of NOBIAS, the max state number of vbSPT, and the starting number of states in SMUAG were all set to 10.

| | **Number of States** | **Running time (s)** | **State 1 ($\mu m^2/s$) (%)** | **State 2 ($\mu m^2/s$) (%)** | **State 3 ($\mu m^2/s$) (%)** | **Transition Matrix** | **Reference** |
|---|---|---|---|---|---|---|---|
| NOBIAS | 3 | 28790.8 | $D_x$: 0.013<br>$D_y$: 0.015<br>56.94 | $D_x$: 0.043<br>$D_y$: 0.049<br>41.29 | $D_x$: 0.675<br>$D_y$: 0.450<br>1.77 | $\begin{bmatrix} 0.933 & 0.067 & 0 \\ 0.098 & 0.897 & 0.005 \\ 0.006 & 0.148 & 0.846 \end{bmatrix}$ | current |
| vbSPT | 10 | 10067.7 | † | † | † | † | Persson 2013 |
| SMAUG | 4 | 264930.6 | 0.0025<br>33.1 | 0.0040<br>49.2 | 0.4121<br>1.4<br><br>* | $\begin{bmatrix} 0.92 & 0.08 & 0 & 0 \\ 0.05 & 0.88 & 0.07 & 0 \\ 0.01 & 0.20 & 0.79 & 0 \\ 0.01 & 0.02 & 0.03 & 0.94 \end{bmatrix}$ | Karslake 2020 |

† The vbSPT suggested a best model of 10 states.

* The SMAUG analysis indicated a 4$^{th}$ state with $D$ = 0.0069 $\mu m^2/s$ and weight fraction = 16.3%.

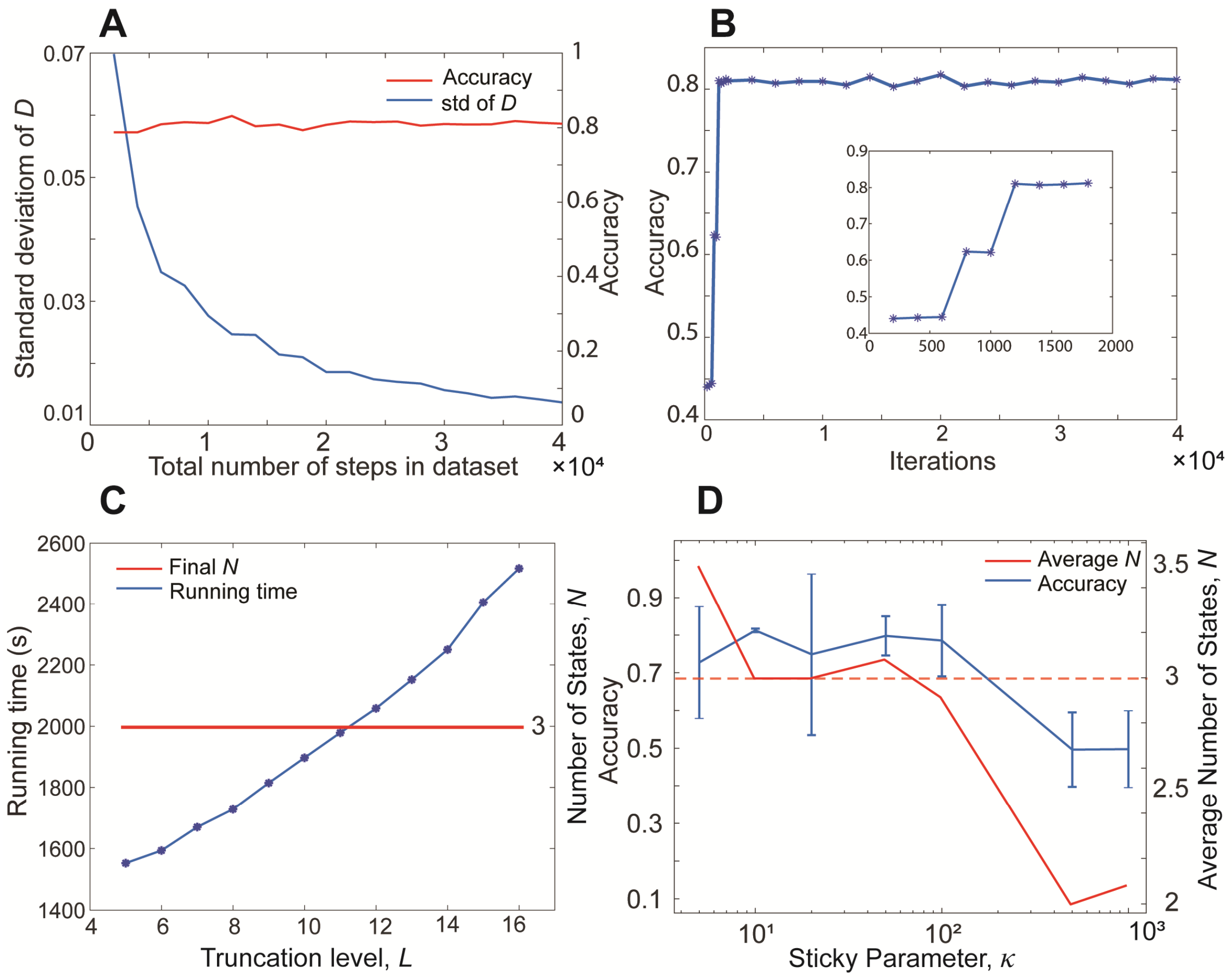


**Figure S1.** Performance evaluation on simulated data. All evaluations use the simulated 3-state motion blur sparse data with the parameter settings as in Table S6, aside from S4C which uses the standard abundant 3-state data. (**A**) The state label accuracy (red) is largely insensitive to the total number of steps in the SPT trajectories, while the posterior parameter sample uncertainty (the standard deviation of the diffusion coefficient, $D$, for the fastest state) improves with an increase in the amount of data amount. All tracks used for this plot are 10 steps long; the data amount was increased by increasing the number of tracks. (**B**) For the same 3-state motion blur sparse dataset, the NOBIAS module accuracy is independent of the final number of iterations beyond 2000 iterations. Inset: zoom in on iterations 0 – 2000. (**C**) The running time (blue) increases with the truncation level, $L$. where the final number of states (red) is not affected. (**D**) Tuning the sticky parameter, $\kappa$, affects the NOBIAS HDP-HMM module performance. Red solid line: average final number of states. The red dashed line indicates the true number of states. Blue line: average state label accuracy (error bars: standard deviation of accuracy over the 12 chains). All results are averaged over 12 chains.

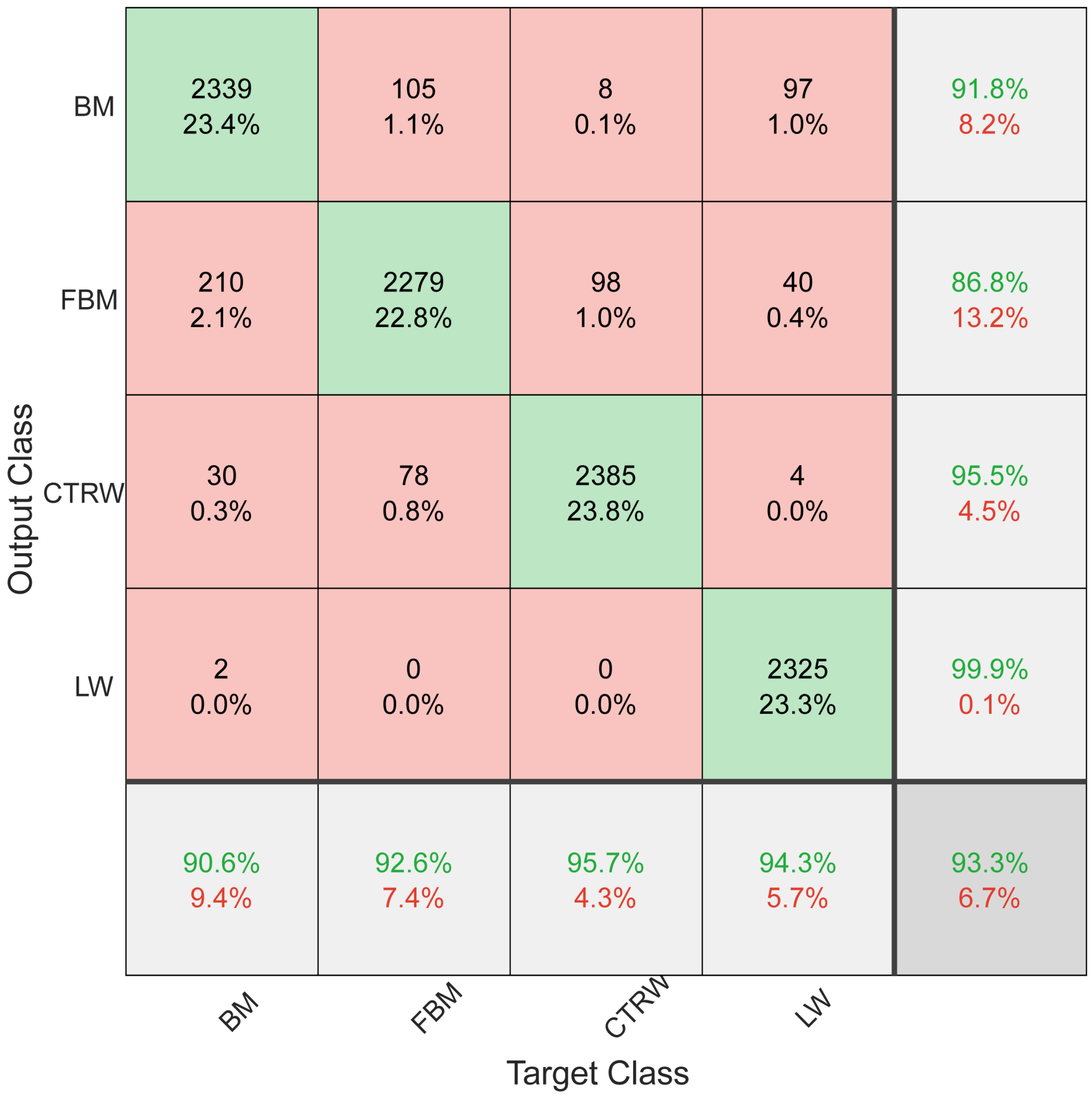


**Figure S2.** Confusion matrix for classification of the diffusion type by the NOBIAS RNN module. A total of 750,000 40-step tracks of the four diffusion types were used to train the network, and 10,000 tracks were tested to get the confusion matrix.

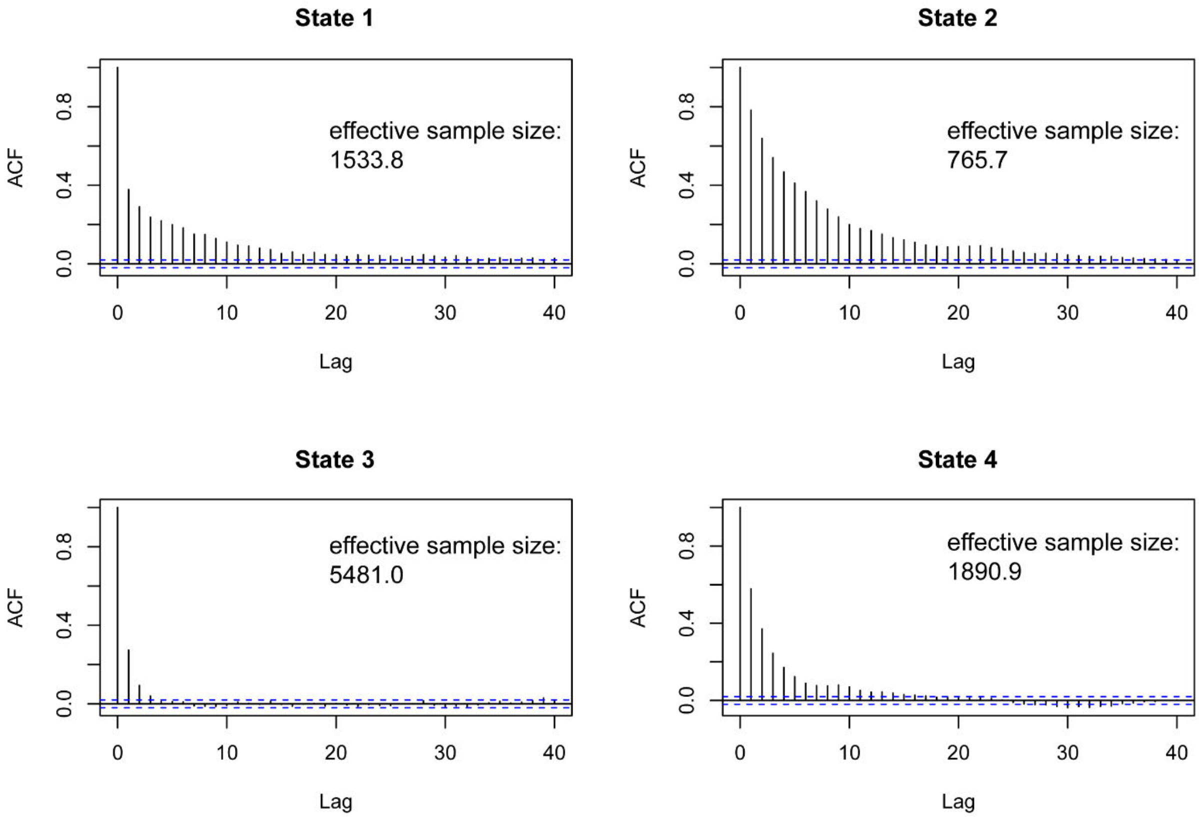


**Figure S3.** Autocorrelation function (ACF) analysis for posterior samples of the diffusion coefficient of the four-state standard abundant simulation described in main text Figure 2E. Over 20,000 iterations, the number of states converges to 4 with a 2000 burn-in. The final 10,000 samples are used for further analysis.

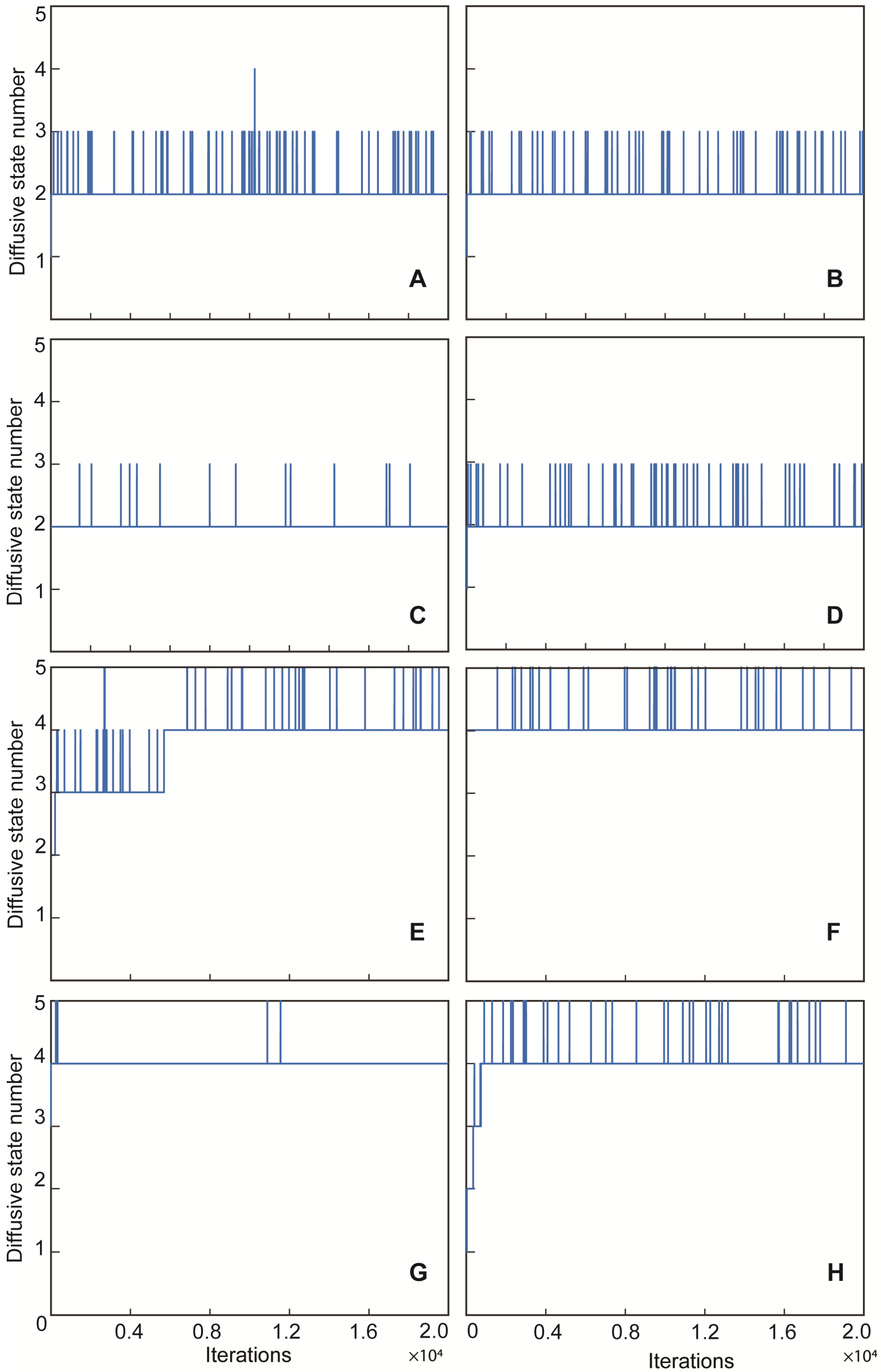


**Figure S4** (continued below)

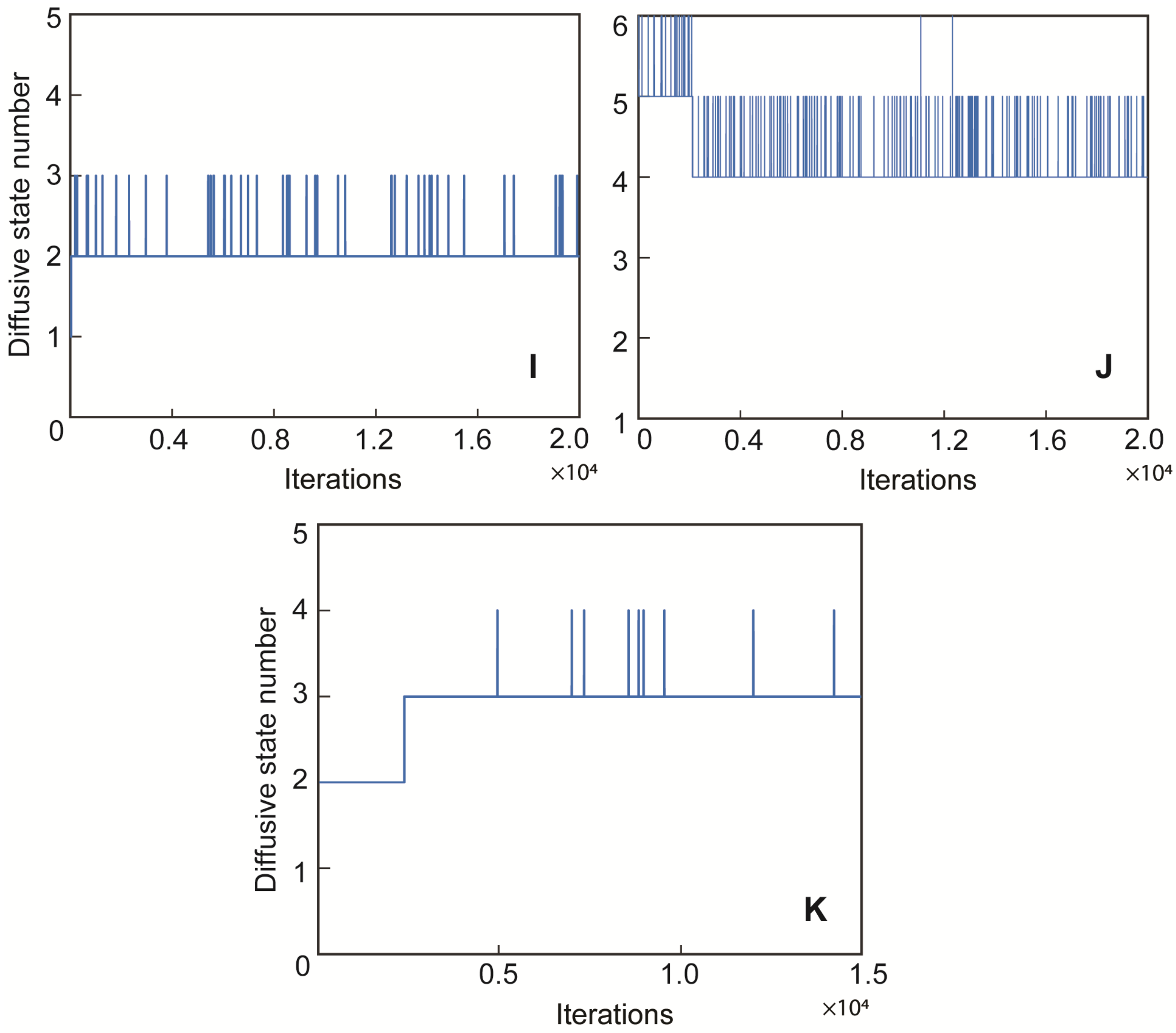


**Figure S4.** Convergence of the number of diffusive states in the NOBIAS HDP-HMM module with iteration number. The number of states convergence plots in A-H correspond to the analysis of simulated tracks in main text Figure 2A-H. The number of states convergence plots in I-J correspond to the analysis of simulated tracks in main text Figure 3A,C. The number of states convergence plot in K corresponds to the analysis of experimental tracks in main text Figure 4B.

**Supplementary Note: Blocked sampler for the NOBIAS HDP-HMM module**

This Blocked sampler for sticky HDP-HMM is mostly based on (Fox, 2009), please find more details and alternative algorithms in the original work. Please see our open source code (https://github.com/BiteenMatlab/NOBIAS) for the implementation of this algorithm in NOBIAS.

The state sequence at the n[th] iteration is sampled as follows based on the state-specific transition probability $\pi^{(n-1)}$ from the last iteration, the global transition distribution $\beta^{(n-1)}$, and the emission parameters $\boldsymbol{\theta}^{(n-1)}$, where $\boldsymbol{\theta} = \{\boldsymbol{\mu},\ \boldsymbol{\Sigma}\}$ include the mean and variance for the multivariate Gaussian:

1. Calculate the backwards message $m_{t+1,t}(k)$, where $k = 1,2, \dots, L$ is the current state. For convenience, note that $\pi = \pi^{(n-1)}$ and $\boldsymbol{\theta} = \boldsymbol{\theta}^{(n-1)}$.
   a. Initialize $m_{T+1,T}(k) = 1$.
   b. For $t = T-1, \dots, 1$ and $k = 1,2, \dots, L$:
$$m_{t,t-1}(k) = \sum_{j=1}^{L} \pi_k(j)\, Norm(\Delta\boldsymbol{x}_t; \mu_j, \Sigma_j) m_{t+1,t}(j)$$
2. Forward sample state assignments sequentially in time. $n_{jk}$ is the transition-counting variable denoting the number of transitions from state $j$ to state $k$, $S_k$ denotes the sufficient statistics for observations in state $k$. Start from $n_{jk} = 0$, $S_k$ is set to empty, and $k, j = 1,2, \dots, L$.
   a. Compute the probability for $\Delta\boldsymbol{x}_t$ in each state $k = 1,2, \dots, L$:
$$p_k(\Delta\boldsymbol{x}_t) = \pi_{z_{t-1}} Norm(\Delta\boldsymbol{x}_t; \mu_k, \Sigma_k) m_{t+1,t}(k)$$
   b. Sample the state assignment $z_t$ at time $t$:
$$z_t \sim \sum_{k=1}^{L} p_k(\Delta\boldsymbol{x}_t) \delta(z_t, k)$$
   c. Add a count to the transition-counting variable and update the cached sufficient statistics for the new assignment $z_t = k$:
$$n_{z_{t-1}z_t} \leftarrow n_{z_{t-1}z_t} + 1 \quad S_k \leftarrow S_k \oplus \Delta\boldsymbol{x}_t$$
   Here, the $\oplus$ operators means add the $\Delta\boldsymbol{x}_t$ to the current cached sufficient statistics.
3. Sample auxiliary variables $m$, $w$, $\bar{m}$ as follows:
   a. For $k, j = 1,2, \dots, L$, and for $n = 1,2, \dots, n_{jk}$, set $m_{jk} = 0$ and sample
$$x \sim \text{Ber}\left(\frac{\alpha\beta_k + \kappa\delta(j,k)}{n + \alpha\beta_k + \kappa\delta(j,k)}\right)$$

With $n$ increments, if $x = 1$, increment $m_{jk}$ by 1.

b. For $j = 1,2, \dots, L$, set $\rho = \kappa/(\kappa + a)$, and sample:

$$w_j \sim \text{Binomial}\left(m_{jj}, \frac{\rho}{\rho + \beta_j(1-\rho)}\right)$$

$$\bar{m}_{jk} = \begin{cases} m_{jk} & j \neq k \\ m_{jj} - w_j & j = k \end{cases}$$

**4.** Update global transition distribution:

$$\beta \sim Dir(\gamma/L + \bar{m}_{\cdot 1}, \dots, \gamma/L + \bar{m}_{\cdot L})$$

**5.** Update the new transition distribution and emission parameters:

$$\pi_k \sim Dir(a\beta_1 + n_{k1}, \dots, a\beta_L + \kappa + n_{kL}, \dots, a\beta_L + n_{kL})$$

$$\theta_k \sim NIW(\theta | \lambda, S_k)$$

**6.** Set $\boldsymbol{\theta}^{(n)} = \boldsymbol{\theta},\ \pi^{(n)} = \pi, \beta^{(n)} = \beta.$
